# Deeply sequenced infectious clones of key cassava begomovirus isolates from Cameroon

**DOI:** 10.1101/2020.08.10.244335

**Authors:** J. Steen Hoyer, Vincent N. Fondong, Mary M. Dallas, Catherine Doyle Aimone, David O. Deppong, Siobain Duffy, Linda Hanley-Bowdoin

## Abstract

We deeply sequenced two pairs of widely used infectious clones (4 plasmids) of the bipartite begomoviruses African cassava mosaic virus (ACMV) and East African cassava mosaic Cameroon virus (EACMCV). The sequences of the ACMV clones were quite divergent from our expectations. We have made raw reads, consensus plasmid sequences, and the infectious clones themselves publicly available.

## Main text

Infectious clones are a central tool of molecular virology. Circular single-stranded DNA viruses such as begomoviruses are often cloned in a two-step process to create partial tandem dimers containing two copies of the virus origin-of-replication hairpin. This configuration is thought to enhance infection by facilitating release of monomer virus segment units by the viral Replication protein (Stenger et al., 1991). Cloned isolates of African cassava mosaic virus (ACMV) and East African cassava mosaic Cameroon virus (EACMCV) from Cameroon provided conclusive proof of synergy between two major clades of cassava begomoviruses (Fondong et al., 2000), which was a defining feature of an epidemic of mosaic disease that devastated cassava production in sub-Saharan Africa (Mbewe et al., 2020; Pita et al., 2001). These clones have been used extensively for molecular genetic analysis (Amin et al., 2011; Beyene et al., 2016; Chauhan et al., 2018; Chellappan et al., 2004, 2005a, 2005b; Chowda Reddy et al., 2008, 2009, 2012; Fondong et al., 2007; Kuria et al., 2017; Ndunguru et al., 2016; Patil et al., 2016; Pita et al., 2001; Reyes et al., 2013; Vanitharani et al., 2003, 2004), including analysis of spontaneous mutation (Aimone et al., 2020; Chen et al., 2019; Fondong and Chen, 2011). Herein we describe new sequence resources for these clones, which essentially confirm the sequence of the EACMCV clones and clarify the identity of the ACMV clones.

Complete and accurate plasmid sequences considerably simplify molecular analysis of infectious clones and design of new constructs. To confirm the sequences of the four plasmids listed in Table 1, we grew transformed *E. coli* DH5α cultures overnight at 37 ºC with ampicillin selection and purified each plasmid with the Qiagen Plasmid Maxi kit. Libraries were prepared from Covaris-sheared plasmid DNA in triplicate with the NEBNext Ultra II kit and sequenced on the Illumina NextSeq 500 platform in the 150 bp paired-end read configuration. Data are available as SRA BioProject PRJNA649777.

**Table 1.** GenBank and Addgene identifiers for the four previously described infectious clones and corresponding virus (monomer) sequences. Six complete sequences are described for the first time here, whereas AF112354 and FJ826890 were previously described (Chowda Reddy et al., 2012; Fondong and Chen, 2011; Fondong et al., 2000).

|  | GenBank accession |  | Addgene ID |
| --- | --- | --- | --- |
|  | Plasmid | Virus |  |
| pBluescript II KS(+) **ACMV** DNA-A 1.4mer | MT856193 | MT858793 | 159134 |
| pUC19 **ACMV** DNA-B 1.5mer | MT856194 | MT858794 | 159135 |
| pBluescript II KS(+) **EACMCV** DNA-A 1.6mer | MT856195 | AF112354 | 159136 |
| pSL1180 **EACMCV** DNA-B 1.6mer | MT856192 | FJ826890 | 159137 |

Three of these plasmids were described by Fondong et al. (2000), and the fourth, for EACMCV DNA-B, was described later (Chowda Reddy et al., 2012; Fondong and Chen, 2011) due to greater difficulty in cloning that sequence in the partial-tandem-dimer configuration. These plasmids had not previously been fully sequenced, so we deduced sequence maps based on the restriction sites used, including for partial-tandem-dimer virus segment inserts. Reads were trimmed with CutAdapt 1.16 (Martin, 2011) and aligned to the hypothesized sequences with the Burrows-Wheeler Aligner (BWA-MEM v0.7.13 (Li, 2013)). Variants relative to each reference sequence were identified with samtools v1.8 (Li et al., 2009) and VarScan v2.4.4 (Koboldt et al., 2012). We corrected each plasmid sequence (Table 1) and aligned reads to it a second time.

The EACMCV DNA-A and DNA-B clones had four and two single-nucleotide differences relative to their corresponding GenBank accessions (AF112354.1 and FJ826890.1). We list these differences in the standard coordinate system (relative to the virus nick site) in the order new sequence-old sequence: T139A, G161R [note ambiguity code in AF112354.1], T181TC, and A206AC for DNA-A and T1671G, A2724AT for DNA-B. These differences may reflect errors or they may be true differences resulting from mutations that occurred in *E. coli*.

The consensus sequences of the ACMV clones, by contrast, were 3.1% and 5.8% divergent from the sequences (AF112352.1 and AF112353.1) originally reported by Fondong et al. (2000), as calculated with Sequence Demarcation Tool v1.2 (Muhire et al., 2014). This difference was not entirely unexpected, because of the parallel history of two sets of ACMV clones: infectious partial-tandem-dimer clones were made via restriction-digestion/ligation from sap-inoculated *Nicotiana benthamiana* plants whereas the monomer-segment-unit clones were cloned with PCR from the same original cassava field sample (Fondong et al., 2000). We expect these complete infectious clone sequences will be of great utility to the community, given that many follow-up publications (Amin et al., 2011; Beyene et al., 2016; Chauhan et al., 2018; Chellappan et al., 2004, 2005a, 2005b; Kuria et al., 2017; Ndunguru et al., 2016; Patil et al., 2016; Reyes et al., 2013; Vanitharani et al., 2003, 2004) specifically referenced the related but non-identical monomer sequences (AF112352.1 and AF112353.1).

We obtained deep coverage, over 18,000-fold across all positions for all four plasmids with an average of 157,000-fold coverage (Figure 1). This read depth was consistent across three separate libraries for each plasmid (see Zenodo record 4075362).

**Figure 1.**
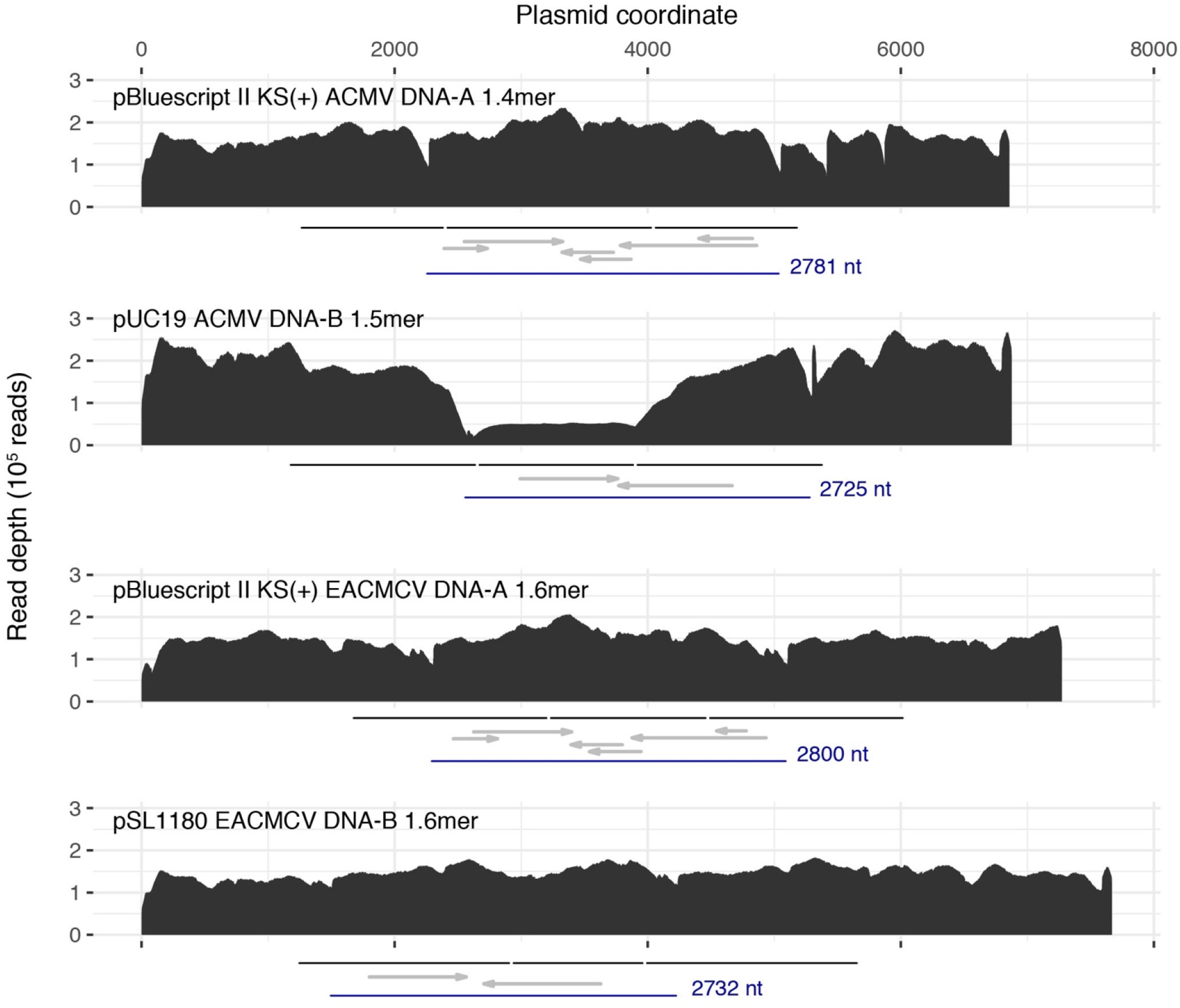
Plots of Illumina read depth across the length of the four infectious clone plasmids. One of three libraries is shown for each plasmid (SRR12354432, SRR12354427, SRR12354424, and SRR12354421). The region in each plasmid corresponding to each virus segment partial-tandem-dimer unit is indicated with a black line under each graph. Vertical white lines demarcate the boundaries of the unique and duplicated regions of each concatemer. Each virus segment monomer unit (in between two replication-origin nick sites) is shown in blue. Canonical virus genes are indicated with gray arrows, left to right for virus sense (AV1, AV2, BV1) and right to left for complementary sense (AC1 to AC4, BC1). Uneven read depth for the ACMV DNA-B plasmid is due to instability (truncation), which is evident in single-cut restriction digests (not shown). Such partial deletion of tandem duplicated regions in *E. coli* is not uncommon (Oliveira et al., 2009).

Our results underscore the value of confirming the sequence of molecular clones. An similar example from the geminivirus literature is the Nigerian Ogoroco clone of ACMV (Briddon et al., 1998).

Kittelmann et al. (Kittelmann et al., 2009) thoroughly documented that the infectious DNA-A clone is not 100% identical to the related GenBank accession (AJ427910.1). We do not believe the large literature on these Cameroonian ACMV isolates requires dramatic reinterpretation, but we recommend using the revised sequences described here for future analyses.

## Data availability

Plasmids are available from Addgene and sequences for full plasmids and ACMV segments are available in GenBank (Table 1). Raw Illumina data are available from the NCBI Sequence Read Archive (PRJNA649777). Data processing code has been archived as Zenodo record 4075362.

## Acknowledgements

This work was supported by NSF award OIA-1545553 to LHB and SD. We thank the NC State University Genomic Sciences Laboratory (Raleigh, NC, USA) for excellent sequencing service. We thank the staff of the Office of Advanced Research Computing (OARC) at Rutgers, The State University of New Jersey, for access to and maintenance of the Amarel cluster. We thank Getu Beyene (Donald Danforth Plant Science Center) for helpful discussions.

